# In addition to being a marker for muscle connective tissue, *Odd skipped-related 2 (OSR2)* is expressed in differentiated muscle cells during chick development

**DOI:** 10.1101/255851

**Authors:** Sonya Nassari, Mickael Orgeur, Cédrine Blavet, Sigmar Stricker, Claire Fournier-Thibault, Delphine Duprez

## Abstract

The zinc finger transcription factor, Odd skipped-related 2 (OSR2) is a recognized marker of connective tissue in chick embryos. OSR2 gain- and loss-of-function experiments indicate a role in irregular connective tissue differentiation in chick limb undifferentiated cells. Re-investigation of *OSR2* transcript location during chick development with in situ hybridization experiments showed that *OSR2* was also expressed in differentiated muscle cells in limbs and head. *OSR2* expression was also observed in differentiated myotubes in chick foetal myoblast cultures. This shows that in addition to being a marker of connective tissue, *OSR2* is also expressed in muscle fibres during chick development.

## INTRODUCTION

Connective tissue (CT) is an important component of the body supporting and connecting organs. During development, reciprocal interactions between CT and myogenic cells are required to form a functional muscular system (reviewed in Nassari et al., 2017a). In limbs, CT cells and myogenic cells are intermingled in muscle masses but have different embryological origins. CT cells originate from lateral plate, while myogenic cells originate from somites (Bourgeois et al., 2015; Chevallier et al., 1977; Kardon, 1998). The muscle differentiation program is under the control of the Myogenic Regulatory Factor (MRFs), including MYF5, MYOD, MRF4 and MYOG (reviewed in Tajbakhsh, 2009). The differentiation program of limb muscle CT cells is not known (reviewed in Nassari et al., 2017a). To date, three types of transcription factors have been identified to regulate muscle formation in a non cell-autonomous manner during limb development: TCF4, a member of the TCF/LEF family (Kardon et al., 2003; Mathew et al., 2011), the T-box transcription factors, TBX4 and TBX5 (Hasson et al., 2010) and the zinc finger transcription factors, Odd skipped-related 1 and 2, OSR1 and OSR2 (Stricker et al., 2012; Vallecillo-García et al., 2017)

The *odd* gene coding for the zinc finger transcription factor odd-skipped was first identified in a screen for gene mutations in *Drosophila* as pair-rule genes involved in body segmentation (Coulter and Wieschaus, 1988; Nüsslein-Volhard and Wieschaus, 1980). In vertebrates, two odd-skipped-related genes, *Osr1* and *Osr2,* have been identified (Lan et al., 2001; So and Danielian, 1999). During mouse and chick embryonic development and organogenesis, *Osr1* and *Osr2* exhibit partially overlapping expression domains in intermediate and lateral mesoderm and at later stages in mesonephros, branchial arches, limbs, mandibular and maxillary prominences (Lan et al., 2001; So and Danielian, 1999; Stricker et al., 2006). *Osr1* and *Osr2* genes display functional equivalence during mouse development, indicating that the distinct functions of *Osr* genes rely on their expression domains (Gao et al., 2009). Consistent with *Osr* expression sites, phenotypes of *Osr* mutant mice indicate a role for *Osr1* in heart and urogenital development (James, 2006; Wang et al., 2005) and a role for *Osr2* in palate and tooth development (Lan et al., 2004; Zhang et al., 2009).

*Osr1* and *Osr2* are recognized markers for irregular CT during chick and mouse development (reviewed in Nassari et al., 2017a). During chick limb development, the expression of both genes labels CT cells and is excluded from myogenic cells, labelled with *PAX3* or *MYOD* (Stricker et al., 2006; Stricker et al., 2012). However, *OSR1/2*-positive cells display partial overlap with tendon *SCX*-positive cells and CT *TCF4*-positive cells in limb buds of embryonic day 4, E4, (HH24) chick embryos. However, *OSRs* are not expressed in tendons when they are formed (Orgeur et al., 2017 bioRxiv posted 20 July 2017 doi:10.1101/165837). Moreover, the overexpression of either OSR1 or OSR2 promotes irregular CT differentiation, while preventing cartilage, tendon and muscle differentiation in chick limb undifferentiated progenitors (Stricker et al., 2012, Orgeur et al., 2017 bioRxiv posted 20 July 2017 doi:10.1101/165837). Conversely, the blockade of OSR1 or OSR2 activity decreases the expression of CT markers, while promoting cartilage marker expression in chick limb undifferentiated progenitors (Stricker et al., 2012). Recently, *Osr1* has been shown to identify a population of embryonic fibro-adipogenic progenitors that has a non cell-autonomous effect on developmental myogenesis (Vallecillo-García et al., 2017). In chick limbs, when the final muscle pattern is set, *OSR2* appears mainly associated with individual muscles, while *OSR1* is expressed in irregular CT within and outside muscles (Nassari et al., 2017b).

In this study, we re-investigated the expression pattern of *OSR2* transcripts during chick development. We showed that in addition to the already known expression in muscle CT, *OSR2* was expressed in myofibres in limbs and head during chick development and in myotubes in muscle cell cultures.

## RESULTS AND DISCUSSION

*OSR2* has been described as being expressed in muscle CT, a subpopulation of irregular CT cells during chick limb development (Nassari et al., 2017b; Stricker et al., 2006; Stricker et al., 2012). We re-investigated *OSR2* expression with in situ hybridization experiments to sections at different developmental stages. We used the chick probe of 361 kb length located between the nucleotides 127 to 487 of *OSR2* gene (Figure 1). This probe has been used previously for gene expression analysis (Stricker et al., 2006; Stricker et al., 2012). Moreover, this OSR2 probe did not crossreact/blast with any gene in chicken genomic databases and did not harbour the zinc finger domain of the *OSR2* gene. At E3, corresponding to HH20 (40 somites), *OSR1* was expressed in limb mesenchyme, while no *OSR2* expression was observed in chick limb buds (Figure 2A,B). *OSR2* was observed in the mesonephros (Figure 2B, arrow), as previously described (Stricker et al., 2006). Muscle progenitors migrate from the hypaxial lips of the dermomyotomes towards the forelimb buds from E2 (HH17, 30 somites) in chick embryos (Chevallier et al., 1977; Tozer et al., 2007). Migrating muscle progenitors assessed with *MYOR* expression did not express *OSR2* at E3/HH20 (Figure 2B,C), showing that *OSR2* was not expressed in muscle progenitors at this stage. At E5 (HH26), the *OSR2* expression domains were similar to that of the myogenic transcription factor *MYOD* in dorsal and ventral limb muscle masses (Figure 2D,E), however it was not clear whether they were expressed in the same cells. From E5 to E8, in addition to being expressed in irregular CT between cartilage elements in distal limb regions and in dermis regions, *OSR2* displayed a strong expression in developing muscles (Nassari et al., 2017b; Stricker et al., 2012). At E9, when the final muscle pattern is set, *OSR2* expression was mostly expressed in muscles regions, in addition to feather buds (Figure 2F). Within muscles, *OSR2* was expressed in a subset of MF20-positive muscle fibres (Figure 2G, arrowheads) and in between MF20-positive cells (Figure 2G, arrows). These results show that in addition to being expressed in irregular and muscle CT during chick limb development, *OSR2* is expressed in differentiated muscle fibres.

**Figure 1.**
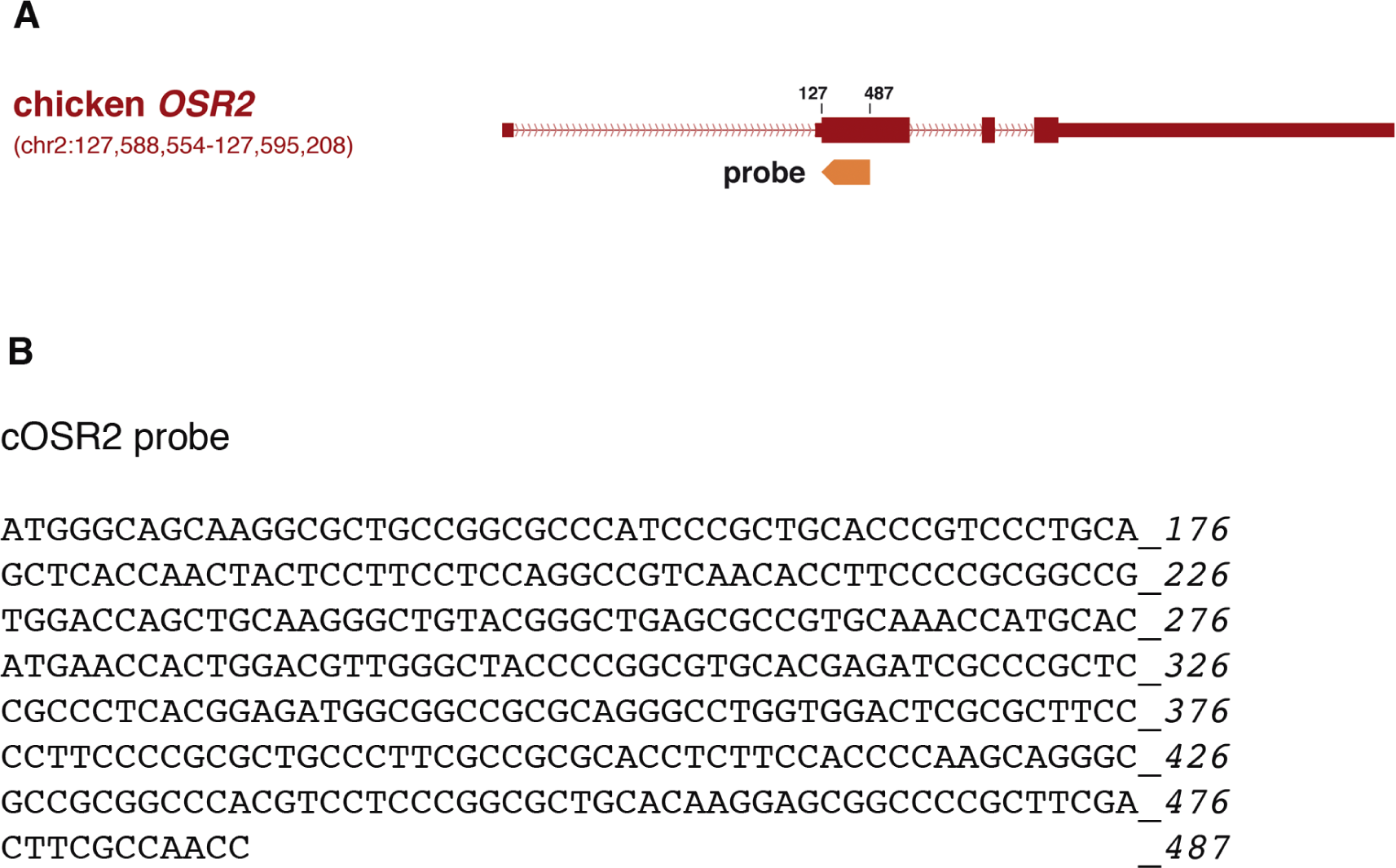
Position of the OSR2 probe along the chicken *OSR2* gene. (**A**) Representation of the *OSR2* gene (NM_001170344.1). Position of the OSR2 probe used for *in situ* hybridization running from nucleotide position 127 to 487 on the *OSR2* gene. (**B**) Sequence of the chick OSR2 probe (361 bp).

**Figure 2.**
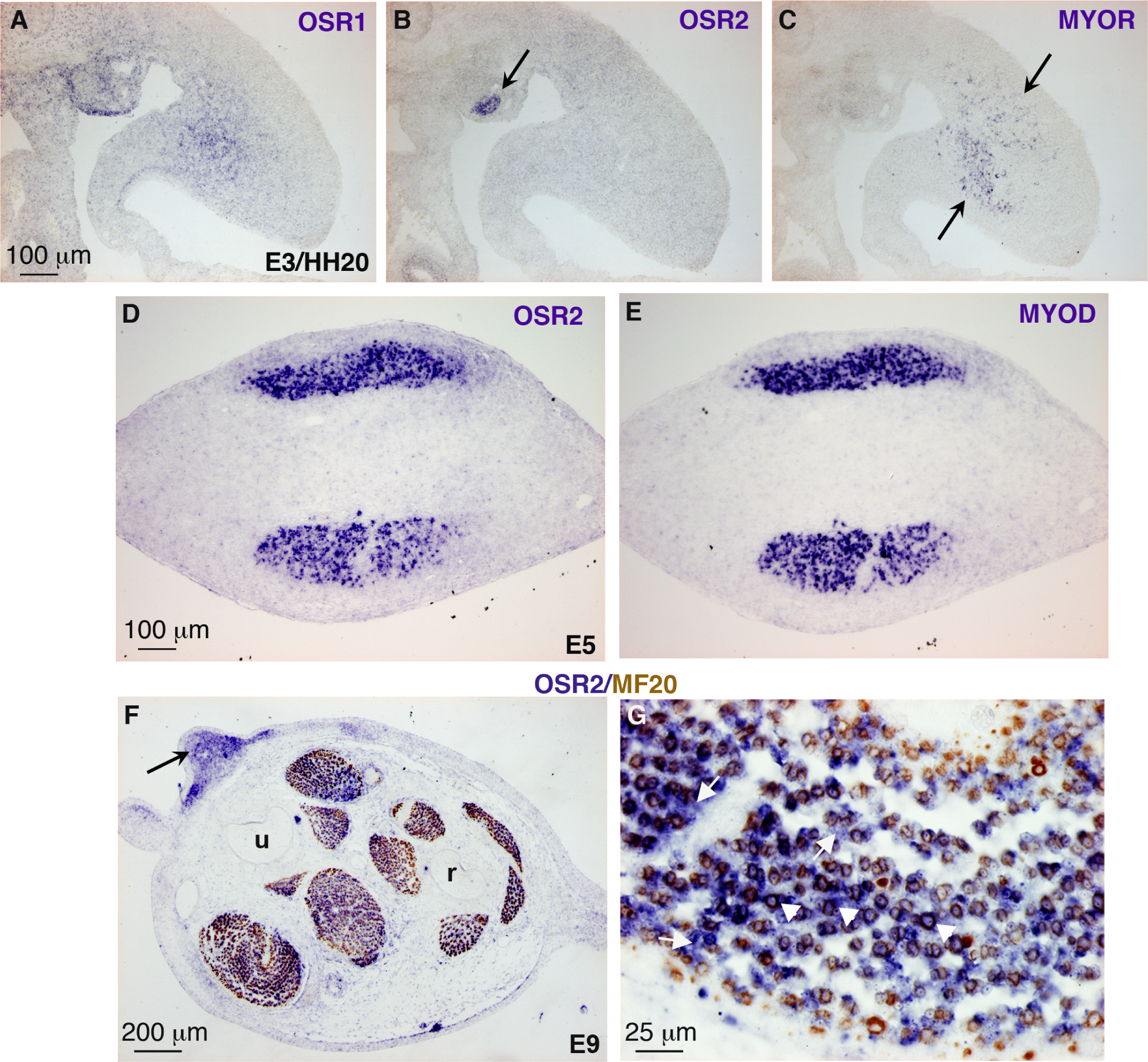
*OSR2* is expressed in CT cells and differentiated muscle cells in limbs. (**A-C**) In situ hybridization to adjacent longitudinal limb sections of E3/HH20 chick embryos with OSR1 (A), OSR2 (B) and MYOR (B) probes (blue). *OSR1* (A) was expressed in limb mesenchyme, while *OSR2* was only detected in mesonephros (B, arrow) and not in muscle progenitors labelled with *MYOR* (C, arrows). (**D,E**) In situ hybridization to adjacent transverse limb sections of E5 chick embryos with OSR2 (D) and MYOD (E) probes (blue). *OSR2* and *MYOD* displayed similar and overlapping expression domains in dorsal and ventral muscle masses. (**F,G**) Transverse forelimb sections of E9 chick embryos were hybridized with OSR2 probe (blue) and then immunostained for myosins using the MF20 antibody (brown). (F) Arrow indicates *OSR2* expression in dermis. (G) Arrowheads indicate double *OSR2*-positive and MF20-positive cells, while arrows point to *OSR2*-positive and MF20-negative cells. r, radius; u, ulna. For all pictures, dorsal is to the top.

Previous studies have highlighted a key role for *OSR2* in the development of specific craniofacial regions (Lan et al., 2004). During craniofacial development of mouse embryos, *Osr2* expression is restricted to specific mesenchymal tissues, including the mesenchyme of the developing palatal shelves and the tongue (Lan et al., 2001). In chick embryos, *OSR2* has been described to be expressed in branchial arches from E3 (Stricker et al., 2006). In order to determine whether *OSR2* was also expressed in differentiated muscle cells during chick craniofacial development, we performed in situ hybridization to head sections of chick embryos at different developmental stages. At E3 (HH20), *OSR1* and *OSR2* were expressed in distinct regions of the first branchial arches outside the muscle regions labelled by *MYOR* (Figure 3A-C). At E4 (HH22), *OSR2* expression was detected in mesenchymal cells of the third branchial arches and did not overlap with that of *MYOR* (Figure 3D-G), which labelled the core of muscle progenitor cells in branchial arches (Grenier et al., 2009). This indicated that *OSR2* was restricted to neural-crest-derived mesenchymal cells in branchial arches before E4. At E7, *OSR2* was strongly expressed in muscle regions in addition to displaying an expression in the head mesenchyme (Figure 3H-J). High magnifications of muscles visualised with transverse (Figure 3K-M) and longitudinal (Figure 3N-P) sections showed that *OSR2* expression was observed in MF20-positive differentiated muscle cells (Figure 3K-P, arrowheads) in addition to muscle CT (Figure 3K-P, arrows).

**Figure 3.**
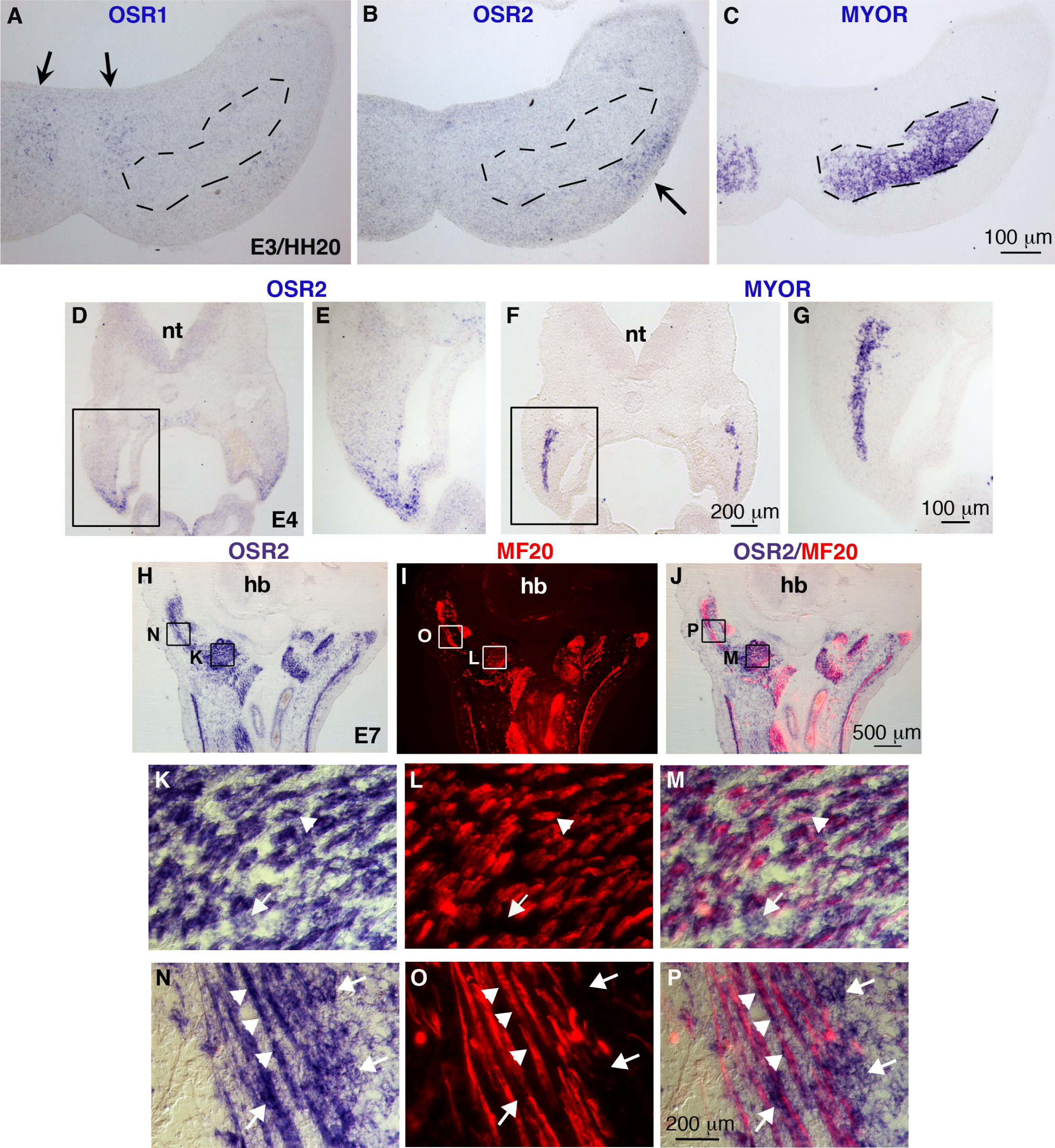
*OSR2* is expressed in CT cells and differentiated muscle cells in branchial arches. (**A-C**) In situ hybridization to adjacent transverse sections of E3/HH20 chick embryos at the level of the first branchial arches (BA1) with OSR1 (A), OSR2 (B) and MYOR (C) probes (blue). (A-C) Arrows point to the *OSR1* and *OSR2* expression in mesenchyme excluded from *MYOR* expression domain that is delineated with dashed lines. (**D-G**) In situ hybridization to adjacent transverse head sections of E4 chick embryos at the level of the third branchial arches (BA3) with OSR2 (D,E) and MYOR (F,G) probes (blue). (E,G) represent high magnifications of framed regions in (D,F), respectively. At E3 and E4, *OSR2* was not detected in the core region of myogenic cells labelled with *MYOR* but was observed in the mesenchyme of the BA1 and BA3. (**H-P**) Sagittal head sections of E7 chick embryos were hybridized with OSR2 probe (blue) and then immunostained for myosins using the MF20 antibody (red). (H-J) 3 panels of the same section are shown: *OSR2* transcripts (H), MF20 labelling (I) and merged *OSR2*/MF20 (J). (K-M) and (N-P) are high magnifications of the framed muscle regions in (H-J), respectively. Arrowheads in (K-P) indicate *OSR2* expression in MF20-positive cells, while arrows point to *OSR2* expression in MF20-negative cells. *OSR2* was detected in both muscle CT and myosin-positive differentiated muscle cells. nt, neural tube; hb, hindbrain.

We also analysed *OSR2* expression *in vitro*, in primary cell cultures of foetal myoblasts isolated from limbs of E10 chicken embryos. Chicken foetal myoblasts were cultured in differentiation culture medium for 2 days. *OSR2* expression was assessed by in situ hybridization (Figure 4A-D) and muscle cell differentiation with MF20 staining (Figure 4A,B,E,F). We observed *OSR2* transcripts in MF20-positive cells (Figure 4A-F, arrows), showing that *OSR2* was expressed in myotubes in chick foetal myoblast cultures

**Figure 4.**
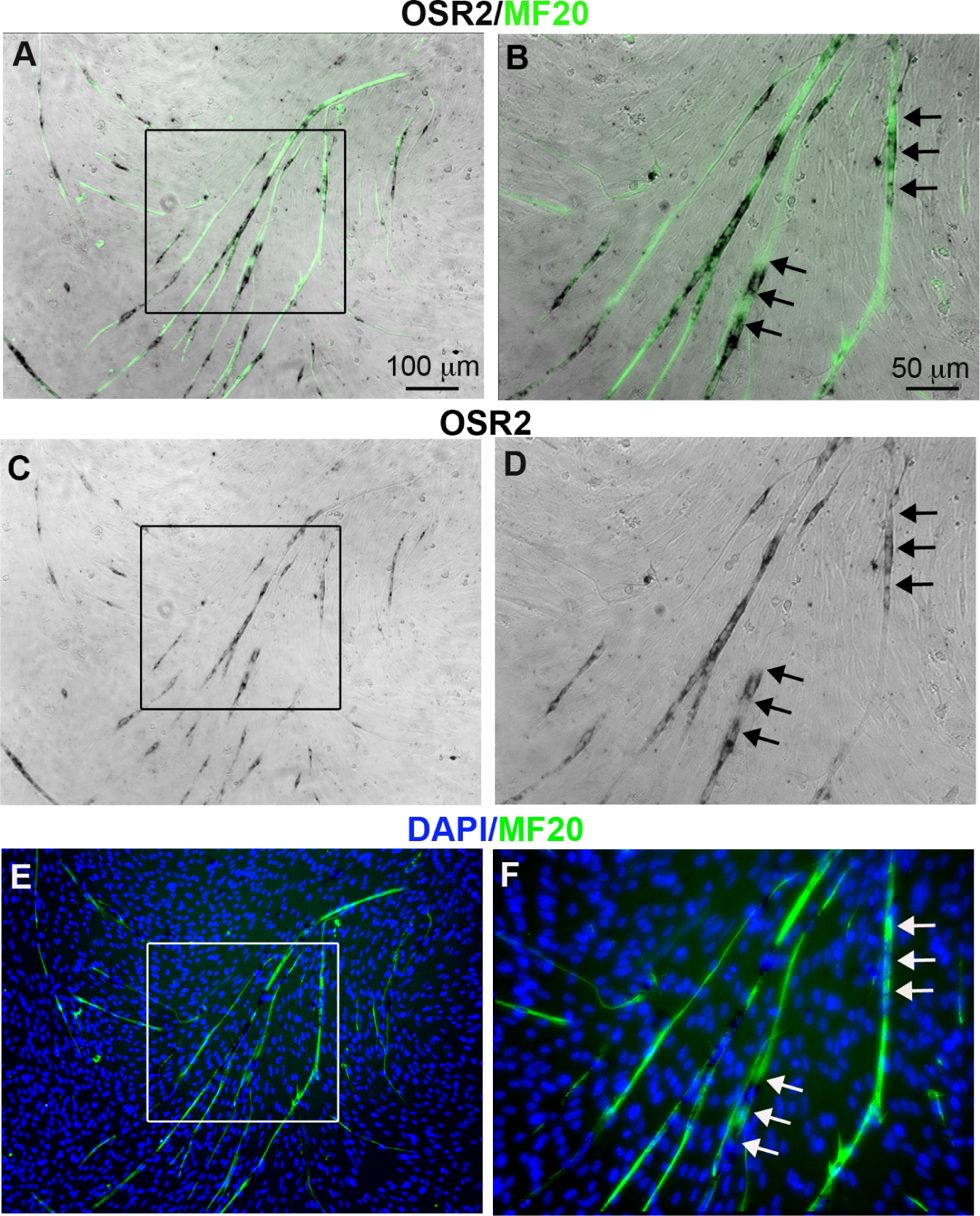
*OSR2* is expressed in differentiated muscle cells *in vitro*. (**A-F**) Chick foetal myoblasts cultured in differentiation conditions were hybridized with *OSR2* probe (black) and immunostained for myosins using the MF20 antibody (green). (A,B) merged of *OSR2* expression and MF20 labelling, (C,D) *OSR2* transcripts, (E,F) MF20 labelling and DAPI staining. (B,D,F) represent high magnifications of the boxed regions in (A,C,E), respectively. Arrows indicate *OSR2* expression in MF20-positive cells.

In addition to being a marker of CT and to promoting CT differentiation from chick mesenchymal stem cells (Nassari et al., 2017b; Stricker et al., 2006; Stricker et al., 2012, Orgeur et al., 2017 bioRxiv posted 20 July 2017 doi:10.1101/165837), we identified *OSR2* as being also expressed in myosin-positive differentiated muscle cells during development of chick embryos. *OSR2* expression was also observed in myotubes of chick foetal myoblast cultures. This unexpected expression of *OSR2* in chick differentiated muscle fibres has never been described before and could bring new insights for OSR2 function during development. It is not clear whether *Osr2* displays similar muscle expression in mice. X-gal staining of forelimb sections of E13.5 and E15.5 *Osr2*^Lacz/−^ embryos did not show any obvious expression in limb muscles (Gao et al., 2011). Moreover, *Osr2^IresCre^* mice displayed specific expression (of reporter gene) in mandibular mesenchyme and did not display any obvious expression in muscle areas in branchial arches of E12.5 mice (Lan et al., 2007). Since previous expression analyses rely on genetic tools, double labelling experiments are nevertheless required to exclude an *Osr2* expression in myotubes during mouse development.

## MATERIALS AND METHODS

### Chick embryos

Fertilized chick eggs from commercial sources (JA57 strain, Institut de Sélection Animale, Lyon, France) were incubated at 38°C in a humidified incubator until appropriate stages. Embryos were staged according to the number of days *in ovo.* Staging using Hamburger and Hamilton, HH (Hamburger and Hamilton, 1992) or somite numbers was also used for young stages.

### Chick myoblast primary cultures

Primary muscle cell cultures were prepared from skeletal muscles of forelimbs of E10 chick foetuses. Muscle cells were mechanically dissociated and seeded in plastic dishes coated with 0.1% gelatine. Myoblast primary cultures were first incubated in a proliferation medium (2/3 Minimum Essential Eagle Medium (MEM), 1/3 Hanks’ salts medium 199, 10% foetal calf serum, 1% penicillin streptomycin and 1% glutamine). At 80% of confluence, differentiation was induced using a differentiation medium (2/3 Minimum Essential Eagle Medium (MEM), 1/3 Hanks’ salts medium 199, 2% foetal calf serum, 1% penicillin streptomycin and 1% glutamine).

### In situ hybridization to embryo sections and cells

Embryos were fixed in 4% paraformaldehyde PBS solution, rinsed successively in 4% and 15% sucrose solution, embedded in a gelatine-sucrose solution (7.5% gelatine, 15% sucrose, 50% Phosphate buffer), and frozen in chilled isopentane. Cryostat sections of 10 to 14 µm were collected on Superfrost/Plus slides (CML, France). Primary muscle cell cultures were fixed in 4% paraformaldehyde PBS solution and rinsed with PBS. Sections and cell cultures were processed for in situ hybridization as previously described (Escot et al., 2013; Nassari et al., 2017b) using digoxigenin-labelled mRNA probes for chick *OSR1*, chick *OSR2* (Nassari et al., 2017b; Stricker et al., 2006), chick *MYOR* and chick *MYOD* (Grenier et al., 2009; von Scheven et al., 2006).

### Immunohistochemistry

Differentiated muscle cells were detected on limb and head sections and chick foetal myoblast cultures after in situ hybridization using the monoclonal antibody against sarcomeric myosin heavy chain, MF20 (Developmental Hybridoma Bank, non-diluted supernatant). Immunohistochemistry was processed after in situ hybridization, as previously described (Grenier et al., 2009; Tozer et al., 2007).

### Image capturing

Images of labelled sections and cultured cells were obtained using a Leica DMI6000 B microscope or a Nikon microscope equipped for epifluorescence. Images were processed using Adobe Photoshop software.

